# Maternal age effects on offspring lifespan and reproduction vary within a species

**DOI:** 10.1101/2023.02.27.530305

**Authors:** Alyssa Liguori, Sovannarith Korm, Alex Profetto, Emily Richters, Kristin E. Gribble

**Author notes:** Corresponding authors: Alyssa Liguori and Kristin Gribble.

## Abstract

Across diverse taxa, offspring from older mothers have decreased lifespan and fitness. Little is known about whether such maternal age effects vary among genotypes for a given species, however. We compared maternal age effects among four strains of rotifers in the *Brachionus plicatilis* species complex. For each strain, we measured lifespan, reproductive schedule, and lifetime reproductive output of offspring produced by young, middle-aged, and old mothers. We found unexpected variability among strains in the magnitude and direction of maternal age effects on offspring life history traits. In one strain, offspring of young mothers lived 20% longer than offspring of old mothers, whereas there were no significant effects of maternal age on lifespan for the other strains. Across strains, advanced maternal age had positive effects, negative effects, or no effect on lifetime reproductive output. For all but one strain, older mothers produced offspring that had higher maximum daily reproduction early in life. Maternal age effects appear to be genetically determined traits, not features of life history strategy or due to accumulation of age-related damage in the germline. Investigating intraspecific variability is critical for understanding the ubiquity of maternal age effects and their role in the evolution of life history and aging.

## 1. Introduction

Maternal age effects, in which a mother’s age at the time of reproduction influences the phenotype of her offspring absent of any changes in genotype, are a common form of intergenerational plasticity. In many taxa, offspring from older mothers have decreased lifespan, fecundity, and health [1–6]. As one of the earliest experimental studies of maternal age effects was conducted by Albert Lansing using the rotifer *Philodina citrina* [3,7], a decrease in lifespan and fecundity in old-mother offspring is often referred to as a “Lansing Effect.” In other species, however, advanced maternal age has positive effects [8–10] or no effects [11] on offspring fitness. While many studies have described maternal age effects, it is unclear how they are distributed across species and whether the tendency to have positive, neutral, or negative maternal age effects is clade-specific. A recent meta-analysis of 97 animal species has shown that Lepidopterans, other invertebrates (including *C. elegans*, rotifers, copepods, annelids, snails, and fruit flies), wild (non-agricultural) mammals, and humans exhibited significant negative effects of advanced maternal age, whereas wild birds had positive maternal age effects on the early development of offspring [12].

The causes of maternal age effects are not clear, but are frequently thought to involve age-related shifts in trade-offs between lifespan and reproduction. If more resources are allocated to higher reproduction, then fewer resources can be allocated to somatic maintenance, repair, and growth, which could result in decreased lifespan, or vice versa [13– 16]. For example, if offspring from older mothers receive fewer maternally-provisioned lipids, gene transcripts, organelles, metabolites, or other resources, they may be shorter-lived or have altered developmental and reproductive schedules. Changes in early life reproductive schedule are thought to impact late-life reproduction or schedule and lifespan [14,17]. Negative maternal age effects are hypothesized to be caused by age-related declines in the reproductive system or gamete quality, or by other physiological constraints as mothers age [3,18]. Positive maternal age effects could occur in organisms with life histories, morphologies, or physiologies that enable parents to allocate more resources to offspring later in life (e.g., if older parents are larger or more experienced) [10,12]. Both positive and negative maternal age effects could be the result of life history adaptations, such as shifting of reproductive schedules, that optimize population fitness across generations in response to environmental changes [18]. However, there is limited evidence to support these hypotheses. With the currently available studies, it is difficult to disentangle the effects of phylogeny, life history, and environment on maternal age effects among broad taxonomic groups [12].

To address questions about the evolution of life history and the impact of maternal age effects on population dynamics, it will be critical to understand the evolutionary mechanisms underlying intergenerational plasticity. A promising avenue of investigation is to characterize intraspecific variability in maternal age effects. Analyses among species are limited by multiple confounding variables, but within species we can compare populations or strains with relatively similar genetics, morphology, life history strategies, and environments, but distinct vital rates. Studies of *Daphnia* [19,20], *Drosophila* [4,11,21,22], and other insects [23] provide examples of high variability in the magnitude and direction of parental age effects on lifespan, development, fecundity, or embryonic diapause among strains and populations. To understand whether intraspecific variability in maternal age effects is typical, additional experimental studies of other clades are needed.

Monogonont rotifers are an ideal study system for investigating phenotypic plasticity across generations. They are microscopic invertebrate animals that are abundant worldwide and play critical roles in many aquatic food webs as primary consumers and prey. Rotifers in the *Brachionus plicatilis* cryptic species complex have lifespans from 1 – 4 weeks and are easily reared across multiple, age-synchronized generations in the laboratory [24,25]. *Brachionus* spp. are cyclical parthenogens: they reproduce asexually, but in response to environmental cues (dense populations) can switch to a sexual reproduction phase that results in the formation of resting eggs [26]. Therefore, genetic diversity of populations can be manipulated, resulting in experimental cohorts that are isogenic, inbred, or genetically diverse. Phenotypic plasticity and maternal effects in response to inter- and intra-specific interactions, including inducible defenses and the induction of sexual reproduction, have been studied in these rotifers for decades [27,28]. Unlike many short-lived invertebrates, *Brachionus* females do not produce large numbers of small offspring in broods, but instead make a large reproductive investment in each offspring, producing 1-6 large neonates each day throughout the reproductive period, with a lifetime reproductive output of 20-30 offspring. This makes it an interesting species in which to examine the conservation of maternal age effects among strains. *Brachionus* has externally brooded embryos, direct development with no larval stages, and no parental care after neonates hatch, so we can distinguish the effects of metamorphoses and maternal care from other maternal age-related differences in maternal provisioning [24,25].

Maternal age effects are just beginning to be explored in *Brachionus* rotifers. Recent studies on the Russian strain of *B. manjavacas* (BmanRUS), have found that advanced maternal age leads to decreased lifespan and lifetime fecundity in offspring [1,29]. While large differences in lifespan and reproductive plasticity between closely-related *Brachionus* strains in response to environmental stressors have been found [30,31], the variability in maternal age effects among *Brachionus* strains is unknown.

Here, we tested for differences in maternal age effects on offspring lifespan and reproduction among four strains from the *Brachionus plicatilis* species complex. We compared one strain of *B. plicatilis* (BpL1) and three strains of *B. manjavacas*: BmanL5 and BmanRUS (both maintained in constant culture for > 20 years) and BmanRUS-RE, which was newly hatched from five-year-old resting (diapausing) eggs of BmanRUS. We compared BmanRUS and BmanRUS-RE to test whether the life history phenotypes that have been observed in past studies were preserved in the diapausing strain (BmanRUS-RE), which had not been subjected to laboratory selective pressures for the past five years [30,31].

Despite their genetic similarity, these strains differ in their origins, time in culture, vital rates under control laboratory conditions, and responses to environmental stressors [32,33]. Under control conditions (21°C, *ad libitum* food), median lifespan ranged from 10 d for BpL1 to 23.5 d for BmanL5. At lower temperatures (16°C), lifespans of BmanRUS and BmanL5 were extended, but the mean lifespan of BpL1 was unaffected [30]. BmanRUS has the highest tendency for mixis (sexual reproduction), BmanL5 has less, and the BpL1 strain is completely amictic (asexual). Under chronic caloric restriction and intermittent fasting diets, BmanRUS lifespan was significantly extended, but BmanL5 and BpL1 had no change in lifespan [31]. Given the substantial differences in lifespan, reproduction, and responses to environment between these strains of the same and closely related species, we wanted to know if they also differ in the effect of maternal age on offspring phenotype.

In this study, we conducted multi-generation life table experiments to quantify survivorship, lifetime reproductive output, and reproductive schedules for multiple maternal age cohorts for each strain. Since BpL1 is the most distantly related strain with the most disparate vital rates and responses to environmental conditions, we expected that it would have different maternal age effects from the other strains. While BmanRUS and BmanRUS-RE are the most closely related, we hypothesized that the BmanRUS line has undergone laboratory evolution, and therefore could have distinct phenotypes from the BmanRUS-RE strain. With these life table data, we tested for trade-offs between lifespan and reproduction and whether relationships between these factors differed among strains and maternal age cohorts. Although previous work on BmanRUS did not find evidence for maternal age-related lifespan-reproduction trade-offs [1,29], we wanted to test whether trade-offs were lacking across the genetically distinct strains studied here. This work will inform future research on the evolution of variability in maternal age effects within and across clades.

## 2. Materials & Methods

### (a) Rotifer and phytoplankton culture

Each rotifer strain was kept in serial culture and fed the chlorophyte algae *Tetraselmis suecica*. Algae cultures were maintained in 2 L flasks of bubbled f/2 medium [34], minus silica, made with 15 ppt Instant Ocean Sea Salt (Instant Ocean, Blacksburg, VA) in distilled water. Both rotifer and algae cultures were grown at 21°C on a 12:12 h light:dark cycle. Cultures of *T. suecica* were maintained in semi-continuous log phase growth by the removal of 40% of the culture and replacement with f/2 medium every other day throughout the experiments.

### (b) Life table experiments

The BmanRUS experiment took place in March 2021. The other three strains were studied simultaneously in July 2021. The same methods were used for all experiments. To control for maternal and grandmaternal ages of the experimental animals, two generations of maternal age synchronization were conducted before the initiation of life table experiments. Amictic eggs were removed from mature females by vortexing, isolated by micropipette, and allowed to hatch and mature for 5 days in *ad libitum* food conditions, at which time eggs were again collected from mature females. After repeating this for two generations, eggs were collected and allowed to hatch for 6 h to initiate the experimental cohort. Thus, the mothers and grandmothers of our experimental F0 cohorts were all 3 – 5 days old.

To initiate and track each cohort, individual neonates were allocated to 1 mL of *T. suecica* at a concentration of 6 × 10^5^ cells mL^-1^ in 15 ppt Instant Ocean in 24-well plates (n = 55 to 75 per strain). Every 24 h, each individual was observed on a Zeiss Stemi 508 microscope, scored as alive or dead, carrying or not carrying eggs (reproductive status), and the number of offspring produced within the previous 24 h was quantified. The original female was then transferred to a new well with new seawater and *T. suecica*. Daily scoring and transfers were conducted until all individuals had died. This method produced individual-level lifespan and reproduction data.

To initiate the F1 generation at young (Y), middle (M), and old (O) maternal ages for each strain, one neonate per mother (hatched within the past 24 h) was pipetted into a well of a new plate with 1 mL of *T. suecica* in Instant Ocean. In some cases, multiple neonates were taken from a single mother, particularly if other mothers had no neonates on the day of collection. These F1 offspring were then tracked for their entire lifetimes in the same manner as their mothers. For BmanL5 (n = 71 - 72) and BmanRUS-RE (n = 60 - 71), experimental F1 generations were initiated at maternal ages of 3 (Y), 6 (M), and 10 d (O). For BpL1, maternal ages were 3 (n = 68), 6 (n =69), and 9 days (n =39) because of its shorter lifespan. The F1 generation of BmanRUS had only two maternal ages: young (3 d; n = 62) and old (11 d; n = 90).

### (c) Statistical analyses

All data were plotted and analyzed using R v. 4.0.2 [35]. The ‘survival’ and ‘survminer’ packages [36,37] were used to create Kaplan-Meier survivorship curves. Differences in survivorship curves among maternal age cohorts were tested within each strain using log-rank/Mantel-Haenszel tests implemented within the ‘survival’ package. For each strain, two separate tests were used to compare Y versus M and Y versus O cohorts (except for BmanRUS, for which there was only Y and O). The Bonferroni method was used to adjust the critical alpha value to account for multiple comparisons. Individuals were right-censored if their death was not observed due to accidental loss during daily transfers.

For lifetime reproductive output (LRO), maximum daily reproduction (MDR), age of 50% LRO, and reproductive period, differences among cohorts were tested within each strain using Kruskal-Wallis rank sum tests. Pairwise comparisons were made using Wilcoxon rank sum tests. To test for life history trade-offs, regression models were fit to describe the relationships between lifespan and LRO, reproductive period (in days and as a percentage of lifespan), and the age of 50% LRO. For each strain, hypothesis tests were conducted to determine whether differences among maternal age cohorts in regression constants and coefficients were statistically significant. These tests were also conducted to compare strains, after pooling maternal age cohorts within each strain.

## 3. Results

### (a) Survivorship

Maternal age effects on offspring lifespan differed among strains (figure 1). For BmanL5, offspring of younger mothers (Y) lived significantly longer than offspring of older mothers (O) (median lifespans of 17 and 13.5 d, respectively; p < 0.0001). Median lifespan of offspring of middle-aged mothers (15 d) was shorter but not significantly different than that of young-mother offspring (p = 0.128). Offspring of younger mothers also lived longer than offspring of older mothers for BmanRUS-RE, but this difference was not significant after a Bonferroni correction for multiple comparisons (median lifespans of 12.5 and 11d, respectively; p = 0.047, Bonferroni corrected alpha = 0.025). For BmanRUS, there was no effect of maternal age on offspring lifespan (p = 0.259); median lifespan for all cohorts was 16 days. There was also no effect of maternal age on offspring lifespan for BpL1 (Y vs. M: p = 0.913; Y vs. O: p = 0.239). Median lifespan for BpL1 cohorts ranged from 8-9 days, much shorter than that for the other strains. Maximum lifespan (age of 5% survivorship) was similar among cohorts for BmanRUS (Y = 21 d, O = 20 d). For BmanL5 and BmanRUS-RE, maximum lifespan was shorter in O cohorts (BmanL5: Y and M = 22 d, O = 20 d; BmanRUS-RE: Y and M = 20 d, O = 17 d). Maximum lifespan was shorter for BpL1 than for other strains across cohorts (Y = 15 d, M = 16 d, O = 14 d; figure 1, table S1).

**Figure 1.**
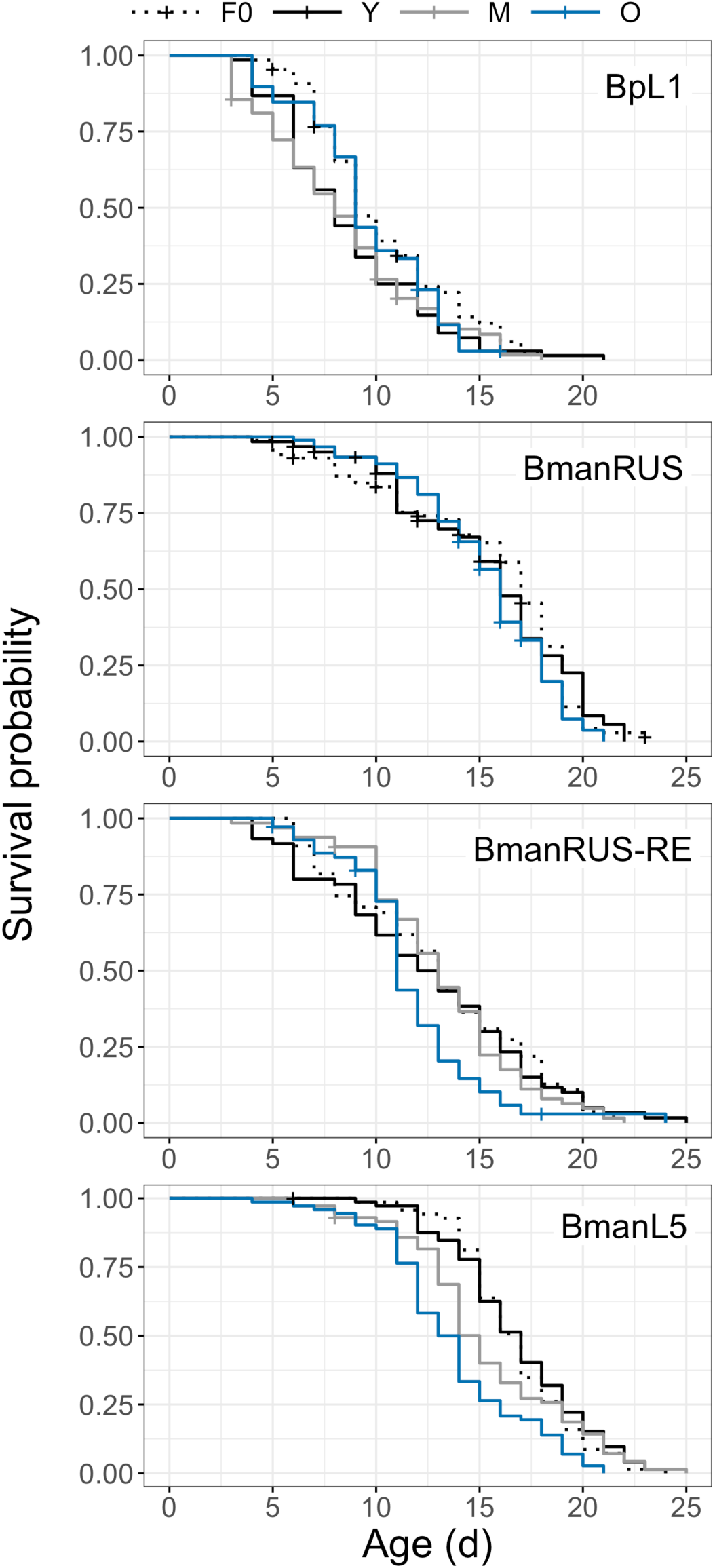
Survivorship curves of the F0 generation (dashed lines) and of the F1 generation from young (Y; black), middle-aged (M; gray), and old (O; blue) mothers, for four *Brachionus* strains.

### (b) Reproduction

Maternal age affected offspring lifetime reproductive output (LRO) in two of four strains, but these effects were in opposing directions (figure 2). For BmanL5, offspring of younger mothers had higher LRO (mean ± SE: 24.8 ± 0.33 neonates ind^-1^) than did offspring of both middle-aged and older mothers (mean ± SE: 22.1 ± 0.46 and 22.2 ± 0.49 neonates ind^-1^, respectively; H(3) = 57.04, p < 0.0001; figure 2G). For BpL1, LRO for all maternal age cohorts was lower than that of the other strains. Offspring of young BpL1 mothers had the lowest LRO (mean ± SE: 8.5 ± 0.61 neonates ind^-1^), compared to older maternal age cohorts that produced just over 10 neonates individual^-1^ on average (H(3) = 9.33, p = 0.025; figure 2A). For both BmanRUS and BmanRUS-RE, there was no significant effect of maternal age on offspring LRO, although offspring from older mothers tended to have higher reproductive output than offspring from younger mothers on average (BmanRUS: H(2) = 2.52, p = 0.28; BmanRUS-RE: H(3) = 8.3, p = 0.04, no significant pairwise comparisons, Wilcoxon rank sum test; figure 2C and 2E, table S1).

**Figure 2.**
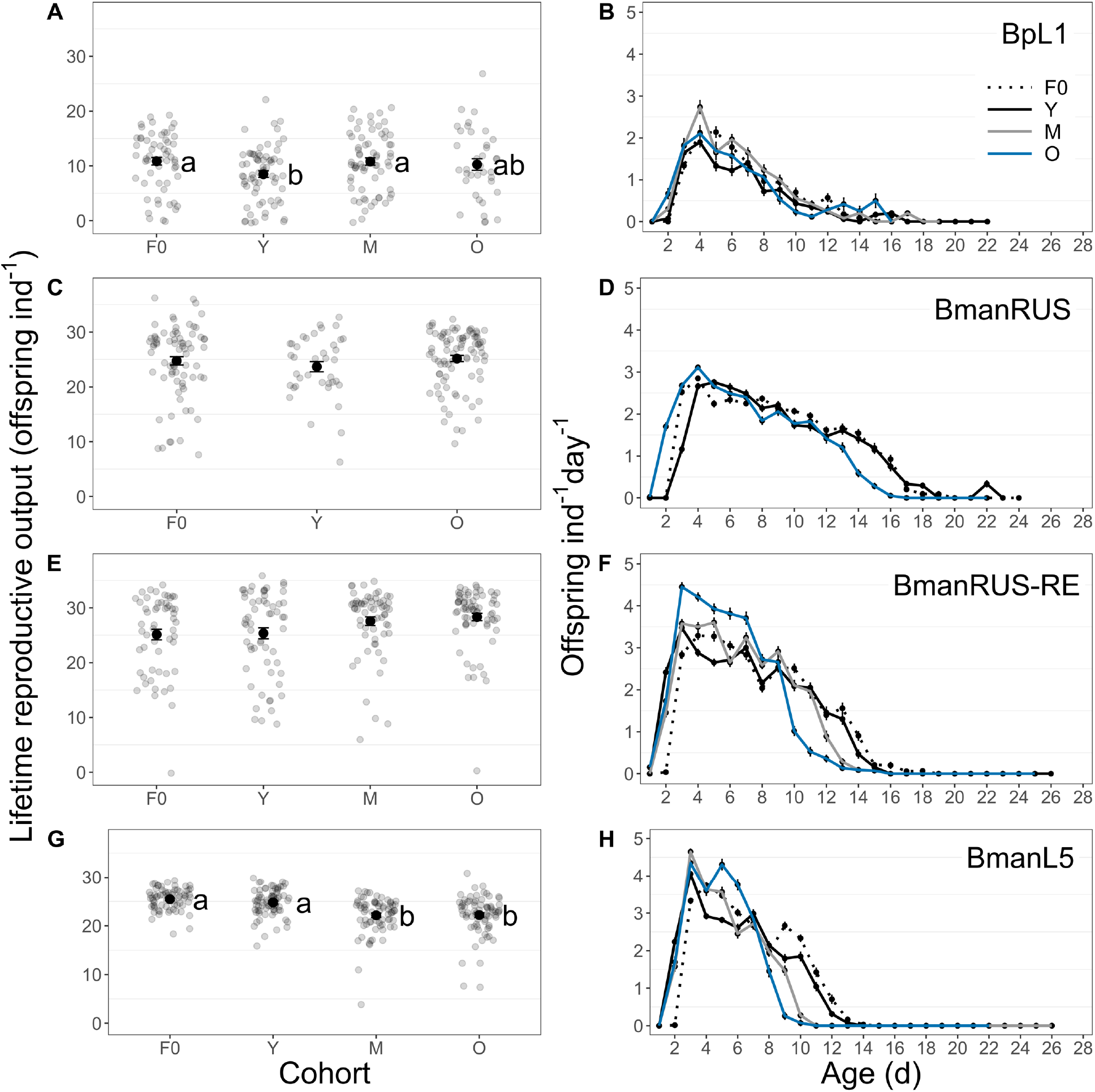
Lifetime reproductive output (LRO; A, C, E, G) and offspring produced per mother per day throughout the lifespan (B, D, F, H), for the BpL1 (A, B), BmanRUS (C, D), BmanRUS-RE (E, F), and BmanL5 (G, H) strains. Mean LRO for the F0 generation and young (Y), middle-aged (M), and old (O) mother cohorts of the F1 generation is shown by bold, black points and individual data points are shown in gray. Significant differences among cohorts are indicated by letters to the right of the mean points (Wilcoxon rank sum tests, alpha = 0.05). Mean offspring production per mother per day is shown by bold points (± SE), and cohorts are indicated by line color (F0 - dashed, Y-black, M - gray, O - blue).

Variability in LRO among individuals also differed among strains and maternal age cohorts. In BpL1, ∼6% of individuals were sterile, which did not occur in the other strains (variance: Y = 24.9, M = 28.9, O = 40.6). In BmanL5, variability in LRO was twice as high in M and O cohorts than in the Y cohort (variance: Y = 7.7, M = 14.6, O = 17.2), with some M and O individuals producing very few offspring (minimum LROs of Y, M, and O cohorts, respectively: 16, 4, and 7 neonates ind^-1^). For BmanRUS-RE, the O cohort had a smaller range in LRO among individuals (17 – 34 neonates ind^-1^, omitting one non-reproductive individual) versus the Y and M cohorts (9 – 36 and 6 – 35 neonates ind^-1^; variance: Y = 58.3, M = 37.8, O = 19.2). Variability in LRO was similar between the Y and O cohorts of BmanRUS (variance: Y = 33.1, O = 27.8; table S1).

Reproductive schedule differed among maternal age cohorts and strains. For the O cohort of BmanL5, daily reproduction was high in early life (mean maximum daily reproduction (MDR): 4.9 neonates ind^-1^; H(3) = 105.4, p < 0.0001), but then declined rapidly after 6 d, until most individuals stopped producing neonates at the age of 9 d. The Y cohort had lower daily reproduction in early life (mean MDR: 4.1 neonates ind^-1^), but also showed a slower decline in reproduction over time. Most Y individuals produced neonates until 12 d old, which led to higher LRO than for M and O cohorts. Rotifers in the M and O cohorts reached 50% LRO one day earlier than rotifers in the Y cohort (age 5 d versus 6 d; H(3) = 156.17, p < 0.0001; figure 2H). A similar pattern was observed for BmanRUS-RE, but the difference in maximum daily reproduction between O and Y cohorts was larger (mean MDR: 4.7 vs. 3.7 neonates ind^-1^; H(3) = 116.15, p < 0.0001). This led to higher LRO in offspring of older mothers on average, despite a steep decline in reproduction that began at day eight. Offspring of young mothers produced at least one neonate per day on average through day 13. Like BmanL5, BmanRUS-RE rotifers in the O cohort reached 50% LRO earlier (age 5 d versus 6 d; H(3) = 38.2, p < 0.0001; figure 2F). For BmanRUS, early peaks in neonate production were not as high as in the other *B. manjavacas* strains. The BmanRUS O cohort peak was higher than the Y peak (mean MDR of O cohort: 3.2 neonates ind^-1^, Y cohort: 3 neonates ind^-1^; H(2) = 16.97, p = 0.0002). After day 13, daily reproduction declined more rapidly in the O cohort than in the Y cohort. Rotifers in the O cohort reached 50% LRO one day earlier than Y cohort rotifers (age 7 d versus 8 d; H(2) = 33.4, p < 0.0001; figure 2D). For BpL1, the highest mean MDR (3 neonates ind^-1^) occurred in the M cohort (H(3) = 22.3, p < 0.0001). There were no significant differences among cohorts in age of 50% LRO (median across cohorts: 4.5 d; H(2) = 1.2, p = 0.55; figure 2B, table S1).

### (c) Reproductive period

Reproductive period, measured as the percent of lifetime in which mothers carried eggs, was affected by maternal age in different ways among strains. For BmanL5 and BmanRUS, Y cohorts had longer reproductive periods than M (BmanL5 only) and O cohorts (BmanL5: H(3) = 32.88, p < 0.0001; BmanRUS: H(2) = 6.04, p = 0.049). Across maternal age cohorts, BmanRUS had longer mean reproductive periods (71.9 to 75.5%) than did BmanL5 (48.8 to 58.6%). There were no significant differences in mean reproductive periods among maternal age cohorts for BmanRUS-RE, although they were longer on average for the Y cohort (73.2 to 78.6%; H(3) = 6.13, p = 0.11). For BpL1, the longest reproductive periods were observed in the M cohort (mean of 65.6% versus 54.6 and 56.8% for the Y and O cohorts; H(3) = 7.77, p = 0.05; figure 3, table S1).

**Figure 3.**
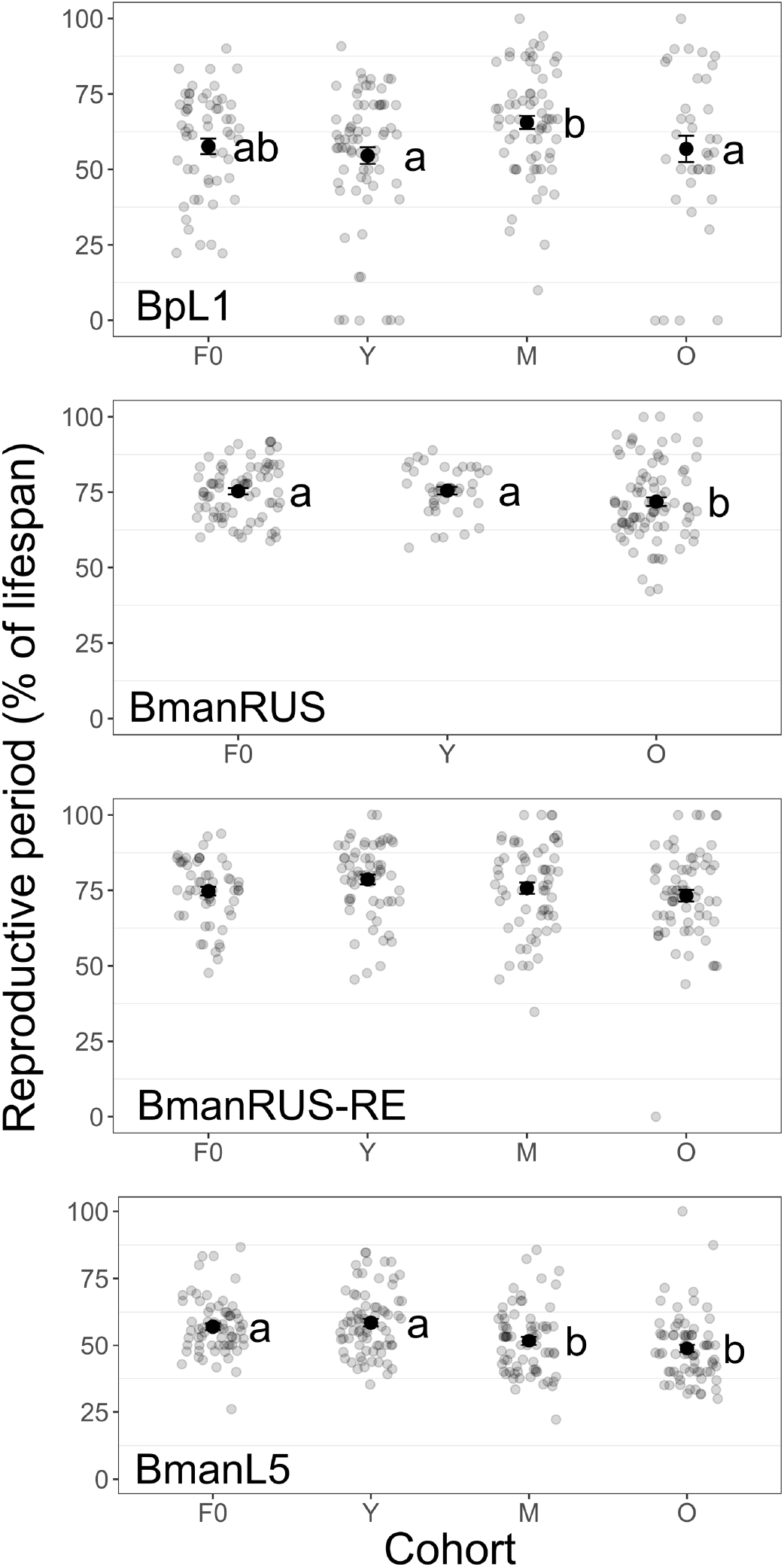
Reproductive period (percent of the lifespan spent carrying eggs) of the F0 generation and young (Y), middle-aged (M), and old (O) mother cohorts of the F1 generation, for four *Brachionus* strains. Mean reproductive periods are shown by bold, black points (± SE) and individual data points are shown in gray. Significant differences among cohorts are indicated by letters to the right of the mean points (Wilcoxon rank sum tests, alpha = 0.05).

### (d) Life history trade-offs

Correlations between lifespan and four metrics of reproduction differed among strains, but there were few differences among maternal age cohorts within strains. LRO increased with lifespan across all strains, but the slope of this relationship was lowest for BmanL5 (slope of 0.39 versus 0.90 - 1.01 for the other strains; figure S1A). For BmanRUS, the relationship between LRO and lifespan was similar between maternal age cohorts (constants: p = 0.13, slopes: p = 0.21; figure 4E). For the remaining strains, only the regression constants (not slopes) differed among maternal age cohorts (BmanL5: p = 0.0009, BmanRUS-RE: p < 0.0001, BpL1: p = 0.0019; figure 4A, 4I, and 4M). The length of the reproductive period (d) also increased with lifespan across strains; BmanL5 had the lowest slope (0.24 versus 0.44 – 0.61 for the other strains; figure S1C). For BmanL5 and BpL1, regression constants differed among maternal age cohorts (BmanL5: p < 0.0001, BpL1: p = 0.03; figure 4O and 4C). For BmanRUS and BmanRUS-RE, the slope of the relationship between reproductive period and lifespan was steeper in the Y cohort versus the older cohorts (BmanRUS slopes: p = 0.009, BmanRUS-RE slopes: p = 0.02; figure 4G and 4K).

**Figure 4.**
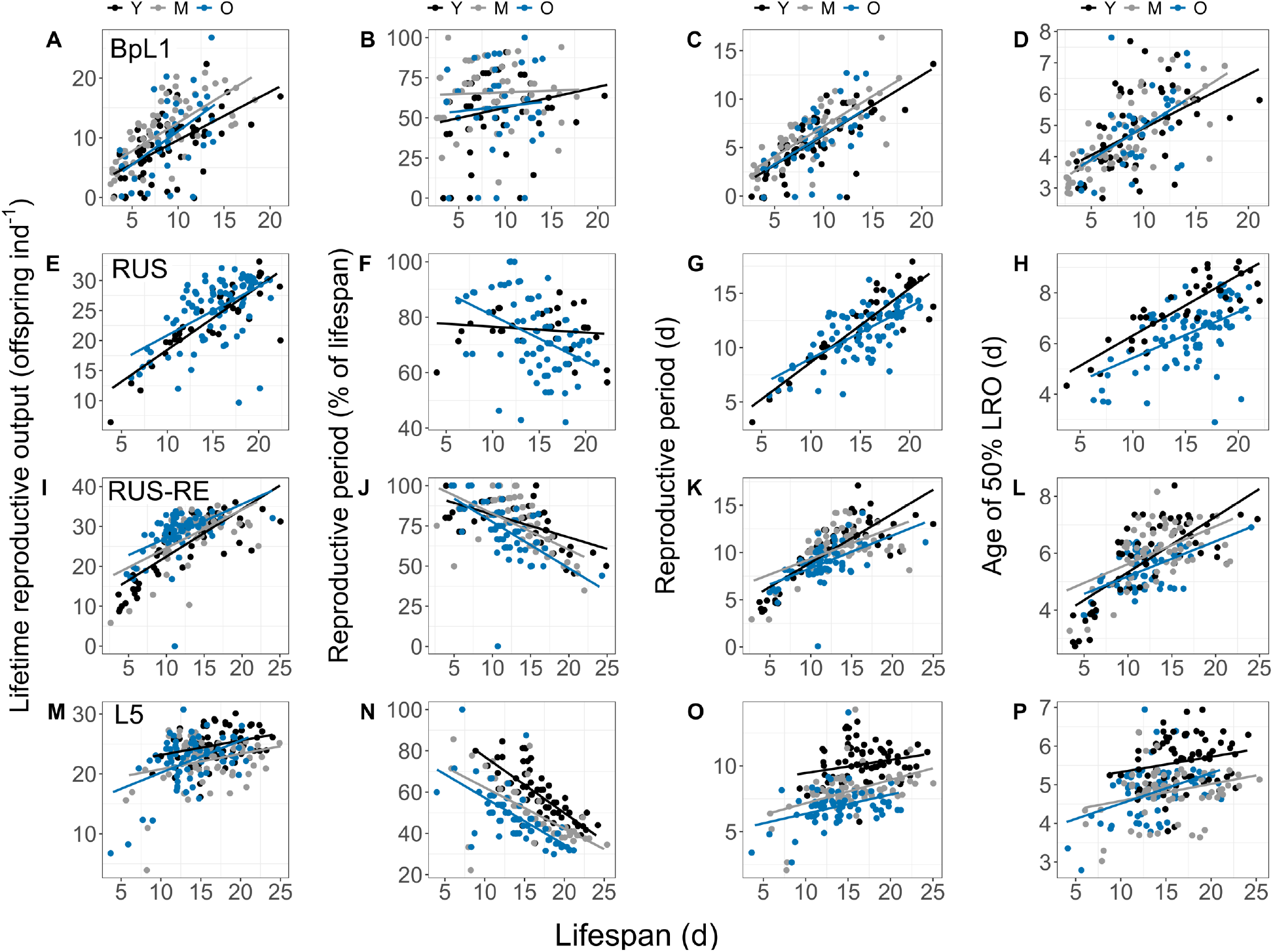
Relationships between lifespan and lifetime reproductive output (LRO; A, E, I, M), reproductive period (as a percent of the lifespan; B, F, J, N), reproductive period in days (C, G, K, O), and age of 50% LRO (D, H, L, P) for the BpL1 (A-D), BmanRUS (E-H), BmanRUS-RE (I-L), and BmanL5 (M-P) strains. Young (Y), middle-aged (M), and old (O) mother cohorts are indicated within each strain by black, gray, and blue, respectively. Note differences between strains in scales of y-axes.

Reproductive period as a percentage of lifespan decreased with increasing lifespan across strains, except for BpL1, for which there was no significant relationship (figure S1B). For BmanRUS and BmanRUS-RE, the negative slope of the relationship was steepest in the O cohorts versus younger cohorts (BmanRUS slopes: p = 0.004, BmanRUS-RE slopes: p = 0.026; figure 4F and 4J). The age of 50% LRO increased with lifespan across strains, but again BmanL5 had the lowest slope (0.076 versus 0.169 – 0.201 for the other strains; figure S1D). For BmanRUS-RE, the slope of this relationship was steepest in the Y cohort (slopes: p = 0.039; figure 4L). Slopes did not differ among maternal age cohorts in the other strains (figure 4D, 4H, and 4P).

## 4. Discussion

Maternal age effects are known to vary among species, in a manner thought to be dependent on large differences in life history strategies among evolutionarily distant taxa. Here we show unexpected intraspecific differences in the magnitude and direction of maternal age effects on offspring lifespan and reproduction in *Brachionus* rotifers. Our findings suggest that maternal age effects are genetically determined traits which may differ even among closely related species with identical life history strategies. This work implies that maternal age effects are not simply caused by the age-related accumulation of cellular or DNA damage that is passed on to offspring. Alternatively, if such damage accumulation does occur, some genotypes may have the capacity to prevent or repair resulting dysfunction in the germline.

Changes in lifespan and reproduction due to maternal age effects were not due to lifespan-reproduction trade-offs. BmanL5, which had the longest post-reproductive period (as a percent of the lifespan), displayed the strongest negative effects of advanced maternal age. Median lifespan of offspring of old mothers was 20% shorter than that of offspring of young mothers. Offspring of young mothers had higher lifetime reproductive output (LRO), with lower variability among individuals, and longer reproductive periods relative to offspring of old mothers.

BmanRUS was least affected by maternal age: both lifespan and LRO were similar between young and old mother cohorts, though advanced maternal age in BmanRUS did lead to shorter reproductive periods and the production of more neonates early in life. For BmanRUS-RE, a strain that was initiated with BmanRUS resting eggs from 2016, offspring of younger mothers had a 10% higher median lifespan (though this difference was not significant), and there was no effect of maternal age on LRO. BmanRUS-RE had the largest difference in early-life reproductive peaks between old and young mother cohorts across strains. For all *B. manjavacas* strains, old mother cohorts produced the majority of their offspring earlier in life than young mother cohorts. For BpL1, the most distantly-related strain with the shortest lifespan and lowest reproduction overall, advanced maternal age led to higher LRO, but had no effect on lifespan.

Intraspecific variability in maternal age effects has been described in only a few other species, the most well-studied of which is *Drosophila melanogaster*. A comparison of six lab strains (4 inbred and 2 outbred) showed that older mothers generally produced offspring with shorter lifespans, but in a single strain, Canton-S, advanced maternal age led to longer lifespan in offspring [22]. We did not observe increased lifespan in old-mother offspring in any rotifer strain in the current study. Lee and colleagues [21] replicated the positive effect of advanced maternal age on offspring lifespan in Canton-S, but saw negative maternal age effects in two additional strains. Bloch Qazi and colleagues [4] also studied Canton-S as well as Oregon-R and found that embryo viability (hatching success) and embryo to adult viability were lower in offspring of old mothers, even in the strain with positive effects of advanced maternal age on offspring lifespan, though the magnitude of effects differed between the strains. Three *D. melanogaster* populations collected from distinct environments in Turkey also showed both positive and negative effects of advanced maternal age on offspring longevity [11]. In *Daphnia pulex*, advanced maternal age decreased offspring lifespan in two of three clones studied, but advanced maternal age led to higher growth rates in offspring in all three clones [19]. In *Daphnia magna*, five clones showed positive, negative, and neutral maternal age effects on lifespan [20]. Such high intraspecific variability in maternal age effects within these model arthropods and rotifers demonstrates the importance of including multiple strains or populations in future studies of maternal age effects in other clades.

Variability in the direction and magnitude of maternal age effects between species has been attributed to differences in life history strategy that result from varied selective pressures and evolutionary constraints. Here, we have shown that strains of the same species can also exhibit divergent maternal age effects. Intraspecific differences in maternal effects could also be due to differential selective pressures in diverse environments. For example, in the striped ground cricket, *Allonemobius fasciatus*, populations that span a wide latitudinal cline face different seasonality and have evolved distinct life histories. Northern populations experiencing short growing seasons only reproduce once in a year (univoltine), while southern populations with longer growing seasons and milder winters can reproduce multiple times per year (bi- and multivoltine). In univoltine populations, maternal age has no effect on the tendency for diapause in offspring. At the end of the short growing season, mating adults produce eggs that diapause, overwinter, and hatch in the spring with sufficient time for maturation. In multivoltine populations, the tendency to produce diapausing eggs increases with maternal age, which is advantageous since the probability of producing offspring that will mature before winter decreases as the growing season progresses [23].

The strains studied here were originally isolated from different geographic sites [31], but all have been continuously cultured in the lab under similar serial culture methods for decades. These strains have been subjected to the same conditions of no predation, a consistent daily light:dark cycle, constant temperature, cycles of high and low food, cycles of high and low population density, and frequent population bottlenecks. Therefore, all strains have been subjected to the same selective pressures for thousands of generations. Differences in initial genetic composition at the time of culture origin, spontaneous mutation, and genetic drift caused by frequent population bottlenecks all may have contributed to the laboratory evolution of these strains, which could underlie the variation in vital rates and maternal age effects observed here.

BmanRUS-RE, which was initiated from resting eggs of BmanRUS collected in 2016, was included in this study to test whether maternal age effect experiment results in BmanRUS from 2014 – 2018 could be replicated [1,29]. We expected that BmanRUS-RE would be genetically and phenotypically more similar to BmanRUS used in previous studies than to the current continuously cultured BmanRUS strain, potentially due to recent laboratory evolution of the BmanRUS strain. In previous BmanRUS studies, offspring of old mothers had shorter lifespans than offspring of young mothers by 10 – 25% and lower LRO by ∼50% [1,29]. Here, maternal age had no effect on lifespan or LRO in BmanRUS. In BmanRUS-RE, there were also no statistically significant effects of maternal age on lifespan or LRO, but on average, offspring from younger mothers lived 1.5 days longer than offspring from old mothers. Thus, the previously observed strong negative maternal age effects in BmanRUS were not fully recovered in BmanRUS-RE. When sexually-produced resting eggs of BmanRUS were collected, stored, and later hatched, it is likely that only a small proportion of the population was randomly sampled. Therefore BmanRUS-RE may be genetically distinct from BmanRUS in both 2016 and 2021, as a result of genetic drift. Such variation within a species suggests that maternal age effects are under genetic control rather than due to a passive, age-related accumulation of damage in the germline or to universal resource trade-offs between reproduction and lifespan.

Life history theory predicts a trade-off between lifespan and reproduction, at least under limiting resources [38]. In the biology of aging literature, this is often interpreted to mean that increased reproduction causes shorter lifespan, or that limiting or delaying reproduction can increase longevity, and that maternal age effects are driven by such trade-offs [13–16]. For example, Plaistow and colleagues [19] attributed the negative lifespan effects of advanced maternal age to earlier reproduction by old mother offspring. In strains of *Daphnia pulex*, they found that advanced maternal age led to shorter lifespan in two of three strains studied. Across all three strains, offspring of older mothers had increased clutch sizes earlier in life and faster growth than offspring of young mothers. Only in the two strains in which lifespan was reduced in offspring of old mothers did reproductive maturation occur at larger body sizes. Thus, the authors concluded that increased early life reproduction traded-off with lifespan and that maternal age effects were a result of better offspring provisioning by older mothers. Similarly, Anderson and colleagues [20] studied multiple strains of *Daphnia magna* and found an “inverse Lansing Effect” in some strains, in which offspring of older mothers lived longer than offspring of younger mothers. They hypothesized that this inverse effect could be a result of *decreased* lipid offspring provisioning by older mothers, which could result in embryonic caloric restriction. The notion of a trade-off between lifespan and reproduction also underlies this hypothesis: decreased lipid stores during development ultimately limits reproduction and subsequently increases lifespan. Although caloric restriction is a well-documented mechanism of lifespan extension in adult animals, however, embryonic and fetal resource limitation has been shown to cause negative outcomes for offspring [39].

In contrast to these other studies, among the *B. manjavacas* strains examined here, earlier or higher reproduction were not uniformly correlated with shorter lifespan. Old-mother cohorts of all three strains produced 50% of their LRO one day earlier, and early life maximum daily reproduction was higher than in young-mother cohorts. BmanRUS-RE had the largest difference between maternal age cohorts in early life reproductive peaks, and BmanRUS had the smallest, but still significant, difference. However, significantly shorter lifespans in the old mother cohort were only observed for BmanL5. Thus, this study does not provide evidence that a trade-off between increased early life reproduction and lifespan drives maternal age effects. Additionally, the effects of maternal age, either positive or negative, could not be explained by trade-offs between reproduction and longevity among the strains. This suggests that trade-offs between somatic maintenance and development time or reproductive output are not primary drivers of lifespan, at least under replete food conditions. This additionally emphasizes the importance of comparing multiple strains before drawing conclusions about the ubiquity of mechanisms controlling life history strategy. Indeed, if we had tested maternal age effects in only BmanL5, in which offspring of older mothers had both shorter lifespans and earlier peaks in reproduction, we might conclude that maternal age effects are driven by life history trade-offs, in a similar manner as the *Daphnia* studies. Conversely, if BmanRUS had been the sole experimental strain, we might eliminate trade-offs between reproduction and lifespan as a potential mechanism, as reproductive timing was affected by maternal age, but lifespan was not.

Maternal age effects on offspring lifespan and reproduction are genotype-specific in *Brachionus* rotifers. Resource trade-offs between lifespan and reproduction likely do not underlie the variable effects observed here. More studies examining intraspecific variability in maternal age effects are needed to determine whether such high variation is widespread among clades or specific to arthropods and rotifers (the most well-studied taxa to date). Future work on maternal age effects in rotifers and other species should include more genotypes, ideally spanning diverse life history strategies. For example, comparing multiple strains with a range of tendencies for mixis and varied vital rates could allow for direct tests of the influence of life history traits on maternal age effects. Future quantitative trait locus (QTL) analyses or comparative genomics studies of strains with varied maternal age effects may provide insight into the genetic controls of intergenerational inheritance.

## Acknowledgements and funding

This work was funded by NSF CAREER Award IOS-1942606 and by NIA R01AG076592 to K. Gribble.

## Data availability

Scripts and raw data are available online: https://github.com/aliguori19/Brachionus_intraspecific_variation

## Supplementary Materials

**Table S1.** Summary statistics for lifespan and reproduction response metrics. Different color cells (white, gray and blue) indicates statistically significant differences among maternal age cohorts within strains. An intermediate shade (light grey) indicates that the cohort is equal to both the white and gray cohorts, which are distinct from each other. Statistical analyses were not conducted for all white columns. (separate Excel file)

**Figure S1.**
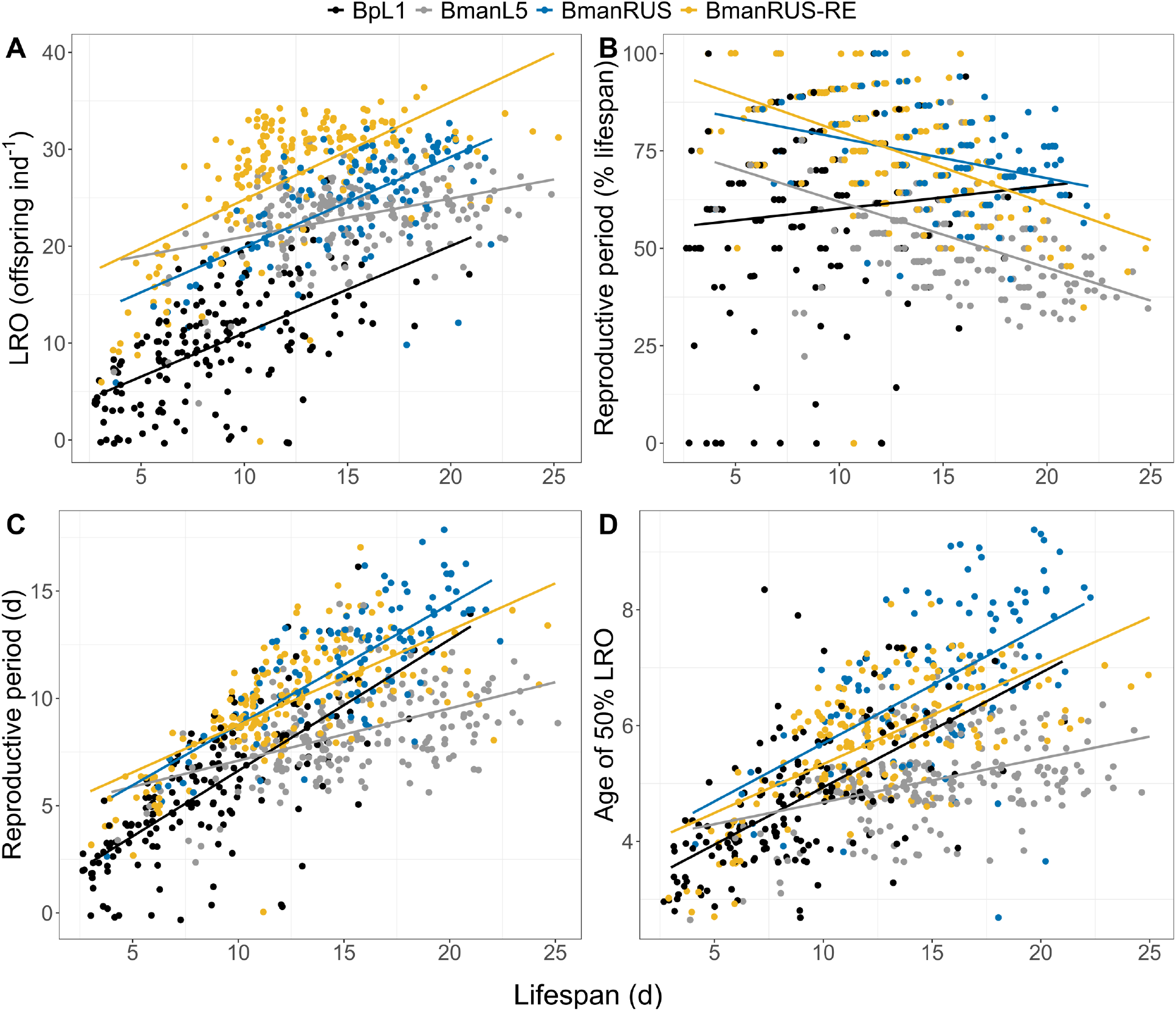
Relationships between lifespan and lifetime reproductive output (LRO; A), reproductive period (as a percent of the lifespan; B), reproductive period in days (C), and age of 50% LRO (D) for the BpL1 (black), BmanL5 (gray), BmanRUS (blue), BmanRUS-RE (goldenrod) strains. Different maternal age cohorts are pooled within each strain here.

## References

1. Bock MJ, Jarvis GC, Corey EL, Stone EE, Gribble KE. 2019 Maternal age alters offspring lifespan, fitness, and lifespan extension under caloric restriction. Sci. Rep. 9, 3138. (doi:10.1038/s41598-019-40011-z)

2. Barclay K, Myrskylä M. 2016 Maternal age and offspring health and health behaviours in late adolescence in Sweden. SSM - Popul. Health 2, 68–76. (doi:10.1016/j.ssmph.2016.02.012)

3. Lansing AI. 1947 A transmissible, cumulative, and reversible factor in aging. J. Gerontol. 2, 228–239. (doi:10.1093/geronj/2.3.228)

4. Bloch Qazi MC, Miller PB, Poeschel PM, Phan MH, Thayer JL, Medrano CL. 2017 Transgenerational effects of maternal and grandmaternal age on offspring viability and performance in Drosophila melanogaster. J. Insect Physiol. 100, 43–52. (doi:10.1016/j.jinsphys.2017.05.007)

5. Beamonte-Barrientos R, Velando A, Drummond H, Torres R. 2010 Senescence of maternal effects: aging influences egg quality and rearing capacities of a long-lived bird. Am. Nat. 175, 469–480. (doi:10.1086/650726)

6. Fox CW, Bush ML, Wallin WG. 2003 Maternal age affects offspring lifespan of the seed beetle, Callosobruchus maculatus. Funct. Ecol. 17, 811–820. (doi:10.1111/j.1365-2435.2003.00799.x)

7. Lansing AI. 1954 A nongenic factor in the longevity of rotifers. Ann. N. Y. Acad. Sci. 57, 455–464. (doi:10.1111/j.1749-6632.1954.tb36418.x)

8. Krishna MS, Santhosh HT, Hegde SN. 2012 Offspring of older males are superior in Drosophila bipectinata. Zool. Stud. 51, 72–84.

9. Perez MF, Francesconi M, Hidalgo-Carcedo C, Lehner B. 2017 Maternal age generates phenotypic variation in Caenorhabditis elegans. Nature 552, 106–109. (doi:10.1038/nature25012)

10. Travers LM, Carlsson H, Lind MI, Maklakov AA. 2021 Beneficial cumulative effects of old parental age on offspring fitness. Proc. R. Soc. B Biol. Sci. 288, 20211843. (doi:10.1098/rspb.2021.1843)

11. Yılmaz M, Özsoy ED, Bozcuk AN. 2008 Maternal age effects on longevity in Drosophila melanogaster populations of different origin. Biogerontology 9, 163–168. (doi:10.1007/s10522-008-9125-y)

12. Ivimey-Cook E, Moorad J. 2020 The diversity of maternal-age effects upon pre-adult survival across animal species. Proc. R. Soc. B Biol. Sci. 287, 20200972. (doi:10.1098/rspb.2020.0972)

13. Stearns SC. 1989 Trade-offs in life-history evolution. Funct. Ecol. 3, 259–268. (doi:10.2307/2389364)

14. Stearns SC. 1976 Life-history tactics: a review of the ideas. Q. Rev. Biol. 51, 3–47. (doi:10.1086/409052)

15. Kirkwood TBL. 1977 Evolution of ageing. Nature 270, 301–304. (doi:10.1038/270301a0)

16. Kirkwood TBL, Holliday R, Maynard Smith J, Holliday R. 1979 The evolution of ageing and longevity. Proc. R. Soc. Lond. B Biol. Sci. 205, 531–546. (doi:10.1098/rspb.1979.0083)

17. Williams GC. 1957 Pleiotropy, natural selection, and the evolution of senescence. Evolution 11, 398–411. (doi:10.2307/2406060)

18. Monaghan P, Maklakov AA, Metcalfe NB. 2020 Intergenerational transfer of ageing: parental age and offspring lifespan. Trends Ecol. Evol. 35, 927–937. (doi:10.1016/j.tree.2020.07.005)

19. Plaistow SJ, Shirley C, Collin H, Cornell SJ, Harney ED. 2015 Offspring provisioning explains clone-specific maternal age effects on life history and life span in the water flea, Daphnia pulex. Am. Nat. 186, 376–389. (doi:10.1086/682277)

20. Anderson CE, Malek MC, Jonas-Closs RA, Cho Y, Peshkin L, Kirschner MW, Yampolsky LY. 2022 Inverse Lansing effect: maternal age and provisioning affecting daughters’ longevity and male offspring production. Am. Nat. (doi:10.1086/721148)

21. Lee J-H, Seo W, Lee S-H, Lee H-Y, Min K-J. 2019 Strain-specific effects of parental age on offspring in Drosophila melanogaster. Entomol. Res. 49, 187–202. (doi:10.1111/1748-5967.12344)

22. Priest NK, Mackowiak B, Promislow DEL. 2002 The role of parental age effects on the evolution of aging. Evolution 56, 927–935. (doi:10.1111/j.0014-3820.2002.tb01405.x)

23. Mousseau TA. 1991 Geographic variation in maternal-age effects on diapause in a cricket. Evolution 45, 1053–1059. (doi:10.2307/2409710)

24. Gribble KE, Snell TW. 2018 Chapter 36 - Rotifers as a Model for the Biology of Aging. In Conn’s Handbook of Models for Human Aging (Second Edition) (eds JL Ram, PM Conn), pp. 483–495. Academic Press. (doi:10.1016/B978-0-12-811353-0.00036-1)

25. Gribble KE. 2021 Brachionus rotifers as a model for investigating dietary and metabolic regulators of aging. Nutr. Healthy Aging 6, 1–15. (doi:10.3233/NHA-200104)

26. Gilbert JJ. 2007 Induction of mictic females in the rotifer Brachionus: oocytes of amictic females respond individually to population-density signal only during oogenesis shortly before oviposition. Freshw. Biol. 52, 1417–1426. (doi:10.1111/j.1365-2427.2007.01782.x)

27. Gilbert JJ. 1966 Rotifer ecology and embryological induction. Science 151, 1234–1237. (doi:10.1126/science.151.3715.1234)

28. Gilbert JJ, McPeek MA. 2013 Maternal age and spine development in a rotifer: ecological implications and evolution. Ecology 94, 2166–2172. (doi:10.1890/13-0768.1)

29. Gribble KE, Jarvis G, Bock M, Mark Welch DB. 2014 Maternal caloric restriction partially rescues the deleterious effects of advanced maternal age on offspring. Aging Cell 13, 623–630. (doi:10.1111/acel.12217)

30. Gribble KE, Moran BM, Jones S, Corey EL, Welch DBM. 2018 Congeneric variability in lifespan extension and onset of senescence suggest active regulation of aging in response to low temperature. Exp. Gerontol. 114, 99–106. (doi:10.1016/j.exger.2018.10.023)

31. Gribble KE, Kaido O, Jarvis G, Mark Welch DB. 2014 Patterns of intraspecific variability in the response to caloric restriction. Exp. Gerontol. 51, 28–37. (doi:10.1016/j.exger.2013.12.005)

32. Mills S, Lunt DH, Gómez A. 2007 Global isolation by distance despite strong regional phylogeography in a small metazoan. BMC Evol. Biol. 7, 225. (doi:10.1186/1471-2148-7-225)

33. Gómez A, Serra M, Carvalho GR, Lunt DH. 2002 Speciation in ancient cryptic species complexes: evidence from the molecular phylogeny of Brachionus plicatilis (Rotifera). Evolution 56, 1431–1444. (doi:10.1111/j.0014-3820.2002.tb01455.x)

34. Guillard RRL. 1975 Culture of Phytoplankton for Feeding Marine Invertebrates. In Culture of Marine Invertebrate Animals: Proceedings — 1st Conference on Culture of Marine Invertebrate Animals Greenport (eds WL Smith, MH Chanley), pp. 29–60. Boston, MA: Springer US. (doi:10.1007/978-1-4615-8714-9_3)

35. R Core Team. 2020 R: A Language and Environment for Statistical Computing. Vienna, Austria: R Foundation for Statistical Computing. See https://www.R-project.org/.

36. Therneau T. 2023 A Package for Survival Analysis in R. See https://CRAN.R-project.org/package=survival.

37. Kassambara A, Kosinski M, Biecek P. 2017 Drawing Survival Curves using ‘ggplot2’. See https://rpkgs.datanovia.com/survminer/index.html.

38. Cohen AA, Coste CFD, Li X-Y, Bourg S, Pavard S. 2020 Are trade-offs really the key drivers of ageing and life span? Funct. Ecol. 34, 153–166. (doi:10.1111/1365-2435.13444)

39. Barker DJ. 1992 The fetal origins of diseases of old age. Eur. J. Clin. Nutr. 46 Suppl 3, S3–9.

